# Extracellular Vesicles promote host immunity during an *M. tuberculosis* infection through RNA Sensing

**DOI:** 10.1101/346254

**Authors:** Yong Cheng, Jeffery S. Schorey

**Affiliations:** Department of Biological Sciences, Eck Institute for Global Health, Center for Rare and Neglected Diseases, University of Notre Dame, Notre Dame, IN 46556.

**Author notes:** Corresponding author. Mailing address: Department of Biology, University of Notre Dame, 130 Galvin Life Science Center, Notre Dame, Indiana, 46556.

**Keywords:** Extracellular Vesicles, *Mycobacterium tuberculosis*, Mycobacterial RNA, SecA2, RIG-I, MAVS, TBK1, IRF3, IFN-β, LC3, Immunotherapy, Macrophages, Mice

## Abstract

Extracellular vesicles (EVs) have been shown to carry microbial components and function in the host defense against infections. In this study, we demonstrate that *Mycobacterium tuberculosis* (*M.tb*) RNA is delivered into macrophage-derived EVs through an *M.tb* SecA2-dependent pathway, and that EVs released from *M.tb*-infected macrophages stimulate a host RIG-I/MAVS/TBK1/IRF3 RNA sensing pathway, leading to type I interferon production in recipient cells. These EVs also promote, in a RIG-I/MAVS-dependent manner, the maturation of *M.tb*-containing phagosomes through a noncanonical LC3 modification, leading to increased bacterial killing. Moreover, treatment of M.tb-infected macrophages or mice with a combination of moxifloxacin and EVs, isolated from *M.tb*-infected macrophages, significantly lowered bacterial burden relative to either treatment alone. We propose that EVs, which are preferentially removed by macrophages *in vivo*, may be developed in combination with effective antibiotics as a novel approach to treat drug-resistant TB.

## Introduction

*Mycobacterium tuberculosis* (*M.tb*), the causative agent of tuberculosis (TB), has been a major instrument of human suffering since antiquity. Presently, over two billion people are infected by *M.tb* worldwide, leading to an estimated 10.4 million active TB cases and 1.7 million deaths in 2016 (WHO report 2017). As an airborne pathogen, *M.tb* primarily infects alveolar macrophages which are exposed to various virulence factors and pathogen-associated molecular patterns (PAMPs). The *M.tb* PAMPs are detected by host germline-encoded pattern-recognition receptors (PRRs) leading to the production of proinflammatory cytokines such as TNF-α and IL-1β, which are essential for an effective immune response (Philips and Ernst, 2012). During an *M.tb* infection, PRR activation also initiates non-transcriptional responses such as the induction of phagocytosis and autophagy in host macrophages (Watson et al., 2012). However, there is limited knowledge on the intercellular trafficking of *M.tb* PAMPs and corresponding activation of the host PRR-dependent pathways in uninfected cells.

Extracellular vesicles (EVs) are membrane-bound vesicles released by both eukaryotic and prokaryotic cells. These vesicles play an important role in intercellular communication regulating various cellular functions of recipient cells. Based on their origin and size, EVs released by eukaryotic cells are divided into three main categories: exosomes, microvesicles and apoptotic bodies (Schorey et al., 2015). Our previous studies found that *M.tb*-infected macrophages release exosomes carrying *M.tb* PAMPs including mycobacterial proteins, lipids and nucleic acids (Bhatnagar et al., 2007; Giri and Schorey, 2010; Singh et al., 2015). These EVs-carrying *M.tb* PAMPs may be detected by PRRs on recipient cells to activate or attenuate cellular responses. For example, EVs from *M.tb*-infected macrophages trigger the TNF-α production in THP-1 human macrophages and naïve mouse bone marrow-derived macrophages (BMMs) (Bhatnagar et al., 2007; Singh et al., 2012). In contrast, these vesicles also suppress the expression of major histocompatibility complex (MHC) class II molecules through a TLR2-dependent pathway in mouse BMMs (Singh et al., 2011). In the context of the adaptive immune response, *M.tb* antigens carried in host cell-derived EVs may be delivered to the antigen processing and presentation pathway in recipient cells. EVs from *Mycobacterium bovis* BCG-infected or *M.tb* culture filtrate protein (CFP)-pulsed macrophages activate an *M.tb* Ag-specific CD4^+^ and CD8^+^ T cells response in naïve mice or mice previously vaccinated with *M. bovis* BCG. The EVs-vaccinated mice were also protected from a low-dose aerosol *M.tb* infection (Giri and Schorey, 2008; Cheng and Schorey; 2013). The recent identification of mycobacterial RNA within EVs (Singh et al., 2015) suggests that host RNA sensors may also be activated in EVs-recipient cells.

In the present study we found that the transport of *M.tb* RNA to EVs is dependent on the expression of the mycobacterial SecA2 secretion system and that EVs carrying *M.tb* RNA stimulate IFN-β production in recipient BMMs. Moreover, EVs also promote LC3-associated *M.tb* phagosome maturation in a RIG-I/MAVS-dependent pathway. Finally, we found EVs from *M.tb*-infected BMMs work synergistically with antibiotics to decrease bacterial load within infected macrophages and following an *in vivo* mouse infection, and do so in a MAVS-dependent manner.

## Results

### EVs Released by *M.tb*-infected Macrophages Stimulate RIG-I/MAVS-dependent Type I Interferon Production in Macrophages

Our previous study identified *M.tb* RNA in EVs isolated from *M.tb*-infected Raw 264.7 cells *in vitro* (Singh et al., 2015). As shown in figure 1A, mycobacterial RNA was also detected in EVs released by mouse BMMs infected with *M.tb*. However, the functional consequence of this EVs-associated mycobacterial RNA in the context of an *M.tb* infection was not defined in this earlier study. During a viral infection, viral RNA is an important PAMP in driving type I IFN production in host cells (Wu and Chen, 2014). Therefore, we hypothesized that EVs-contained *M.tb* RNA may stimulate the host nucleic acid sensing pathways, triggering type I IFN production in recipient cells. As shown in Fig.1B and 1C, EVs secreted from *M.tb*-infected BMMs induced in naive BMMs a dose-dependent production of IFN-β. Maximum IFN-β mRNA transcription was observed 4 hours post treatment when using a concentration of 10 μg/ml EVs isolated from *M.tb*-infected macrophages (Fig.1D). In contrast, no IFN-β mRNA induction was detected in cells treated with EVs from uninfected cells (Fig.1D). MAVS and MyD88 are two critical adaptor proteins in host RNA sensing pathways that perceive cytosolic and endosomal foreigner RNA respectively and drive type I IFN production (Wu and Chen, 2014). To test their roles in EVs-induced type I IFN production, IFN-β expression was measured in MAVS-knockdown and MyD88-deficient BMMs following EV treatment. As shown in Fig. 1E and 1F, when using MAVS-knockdown BMMs as the recipient cells, EVs isolated from *M.tb*-infected macrophages failed to induce IFN-β production. However, no significant difference in IFN-β production was seen between MyD88-deficient and WT BMMs (Fig. S1A and S1B), suggesting a role for the host cytosolic RNA sensing pathway in EVs-induced type I IFN production. Similar, a loss of EVs-induced IFN-β production was also detected in *Mavs* ^−/−^ BMMs (Fig. 1G and 1H). MAVS are activated following interaction with either of two cytosolic RNA sensors RIG-I and MDA5 that recognize foreign RNA (Wu and Chen, 2014). The importance of RIG-I in type I IFN production during a bacterial infection has been assessed in *Listeria monocytogenes* (Abdullah et al., 2012). To test if RIG-I is also involved in EVs-induced type I IFN production, we measured the IFN-β production in RIG-I-knockdown BMMs treated with EVs from *M.tb*-infected macrophages. Similar to the *Mavs* ^−/−^, knockdown of RIG-I in BMMs significantly diminished IFN-β production following treatment with EVs from *M.tb*-infected macrophages compared to control siRNA-treated cells (Fig. 1E, 1F and S1E). In contrast, neither RIG-I or MAVS-knockdown has significant effect on the TNF-α production in BMMs treated with EVs secreted from *M.tb*-infected macrophages (<u>Fig. S1C and S1D</u>). To determine if MAVS were required for IFN-β production by non-infected cells in the presence of infected macrophages, WT *M.tb*-infected BMMs were co-cultured with uninfected WT and *Mavs* ^−/−^ BMMs using a transwell system (Fig. 1I), and IFN-β mRNA abundance in cells of the bottom chamber was measured. In *Mavs* ^−/−^ BMMs, the IFN-β expression was almost undectable, while produced in abundance by WT BMMs (Fig. 1J).

**Figure 1.**
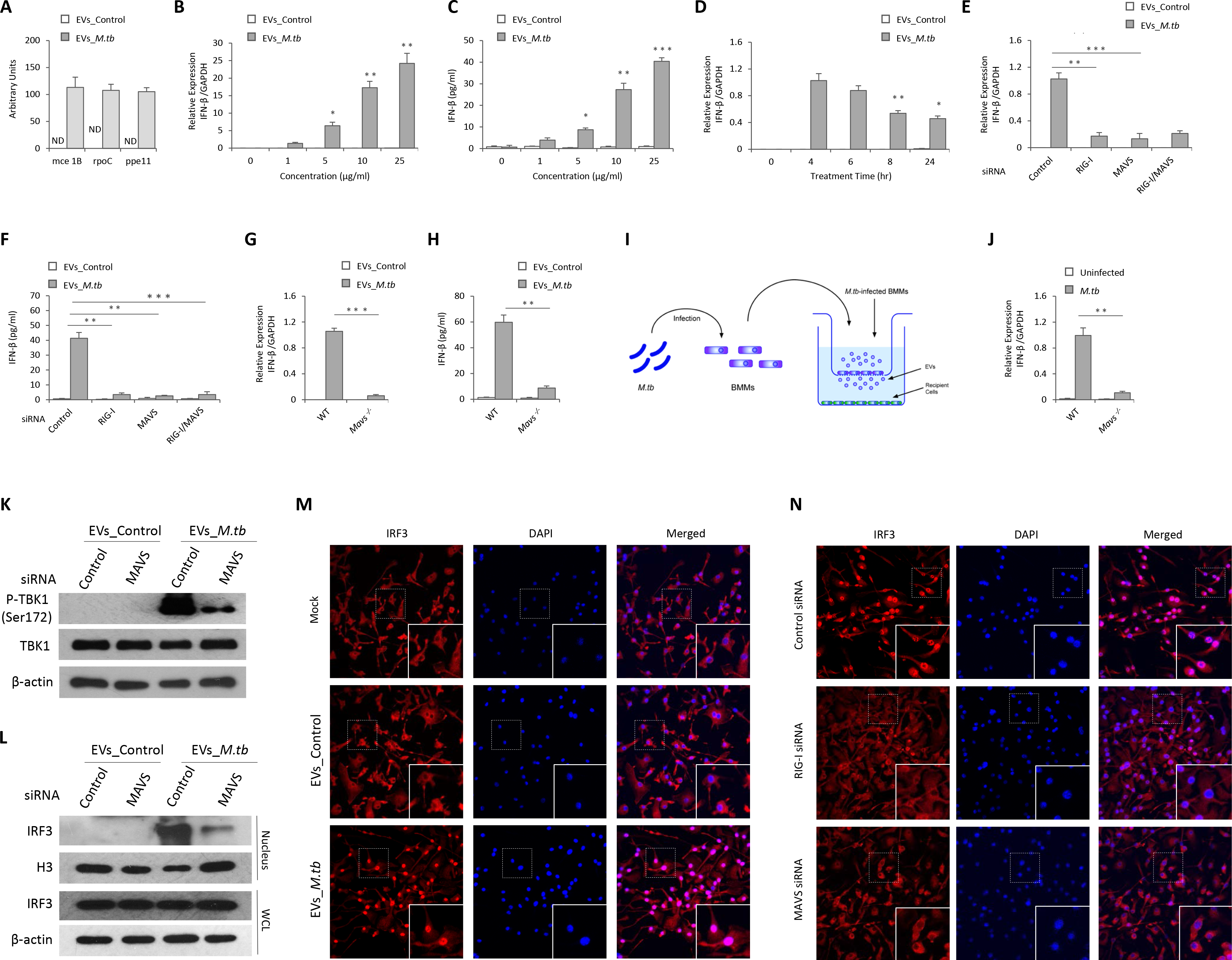
EVs Released by *M.tb*-infected Macrophages Stimulate RIG-I/MAVS-dependent Type I Interferon Expression in Host Cells. **(A)** Quantitative RT-PCR analysis of *M.tb* RNA in EVs from uninfected (EVs_Control) or *M.tb*-infected (EVs_*M.tb*) BMMs. ND, not detected. Quantitative RT-PCR analysis for IFN-β mRNA **(B)** or IFN-β protein **(C)** levels in wild type BMMs 5 and 24 hr post-treatment with EVs respectively. **(D)** IFN-β mRNA level was quantified in wild type BMMs treated with EVs at a concentration of 10μg/ml for the times indicated. Quantitative RT-PCR analysis for IFN-β mRNA **(E)** or IFN-β protein **(F)** levels in either WT, MAVS-and/or RIG-I-knockdown BMMs treated for 4 hr (E) or 24 hr (F) with 10μg/ml EVs from uninfected or *M.tb*-infected macrophages. **(G)** and **(H)** Similar to above except using WT and *Mavs* ^−/−^ BMMs. **(I)** Schematic of transwell assay for measuring IFN-β induction. BMMs were uninfected or infected with WT *M.tb* at a MOI of 5. Twenty-four hr post-infection, cells were transferred into the top chamber and co-cultured with naïve BMMs (Bottom chamber). The IFN-β mRNA levels in BMMs (bottom) were quantified by qRT-PCR after 24 hr. **(J)** Top chamber: WT BMMs; Bottom chamber: WT or *Mavs* ^−/−^ BMMs. **(K)** Western blot analysis for phospho-TBK1 (Ser172) in WT and MAVS-knockdown BMMs treated for 4 hr with EVs from uninfected or *M.tb*-infected macrophages. β-actin served as a loading control. **(L)** Western blot analysis of IRF3 under the conditions described above for TBK1. Histone H3 (H3) and β-actin were used as loading controls for nuclear fraction and whole-cell lysate (WCL) respectively. **(M)** Fluorescence microscopy analysis for nuclear translocation of IRF3 in wild type BMMs untreated (Mock) or treated with EVs from uninfected or *M.tb*-infected macrophages. **(N)** IRF3 nuclear translocation was analyzed in RIG-I-or MAVS-knockdown BMMs. Data shown in (A) - (H), and (J) are the mean ± SD (n = 3 per group) and representative of three independent experiments. * p < 0.05, ** p < 0.01 and *** p < 0.001 by Student’s t-test (two tailed).

### EVs Released by *M.tb*-infected BMMs Activate TBK1 and IRF3 in Macrophages

Protein kinase TBK1 and transcriptional regulator IRF3 are two critical factors downstream of the MAVS-dependent RNA signaling pathway during a viral infection. Following stimulation, IRF3 is phosphorylated by TBK1 and subsequently transported into the nucleus to initiate transcription of type I IFNs (Wu and Chen, 2014). The EVs released by *M.tb*-infected macrophages induced TBK1 phosphorylation at Ser172 as well as IRF3 nuclear translocation (Fig. 1K and 1L). Both TBK1 phosphorylation and IRF3 nuclear translocation was significantly attenuated in MAVS-knockdown BMMs. A similar lack of response was observed in BMMs treated with EVs from uninfected macrophages. Immunofluorescence microscopy analysis also showed the nuclear translocation of IRF3 in WT BMMs following exposure with EVs from *M.tb*-infected macrophages but this translocation was absent when using RIG-I-or MAVS-knockdown BMMs (Fig. 1M and 1N).

### EVs-induced Type I Production in BMMs Requires the *M.tb* SecA2 Secretion System

The SecA2 and ESX-1 protein secretion systems are important for mycobacterial virulence (Feltcher and Braunstein, 2012; Gröschel et al., 2016). Recently, SecA2 was shown to be required for the secretion of *Listeria monocytogenes* nucleic acids while Esx-1 was required for mycobacterial DNA release into the cytosol of infected cells (Abdullah et al., 2012, Manzanillo et al., 2012). To determine whether these secretion systems were also required for *M.tb* RNA trafficking into EVs within infected macrophages, we analyzed EVs released by macrophages infected with a *ΔsecA2* or *ΔesxA M.tb*. As seen in Fig. 2A-2D, the deficiency of either the *secA2* or *esxA* had no significant effect on the EV biogenesis by infected macrophages. EVs isolated from macrophages infected with the *M.tb* strains maintain a similar size profile as those from uninfected macrophages (Fig. 2A and 2C). Additionally, a similar EV yield was achieved across all samples (Fig. 2B and 2D). Quantitative RT-PCR was performed to define the amount of *M.tb* RNA in the isolated EVs. A significant decrease in *M.tb* RNA was seen in EVs released from macrophages infected with the *ΔsecA2 M.tb* when compared to vesicles released from cells infected with WT or *secA2* complementary *M.tb* strains (Fig. 2E). No significant difference in *M.tb* RNA abundance was detected among EVs purified from macrophages infected with WT, Δ*esxA* or *esxA* complementary *M.tb* strains (Fig. 2F). Furthermore, EVs released from macrophages infected with the Δ*secA2 M.tb* failed to induce IFN-β production in recipient BMMs (Fig. 2G and 2H). This lack of IFN-β production by EVs-treated BMMs was rescued by adding liposome-encapsulated RNA using RNA that was isolated from EVs released by WT *M.tb*-infected macrophages (Fig. 2I). No significant difference in IFN-β production was detected in BMMs treated with EVs from macrophages infected with WT, Δ*esxA* or esxA complementary *M.tb* strains (Fig. 2J and 2K).

**Figure 2.**
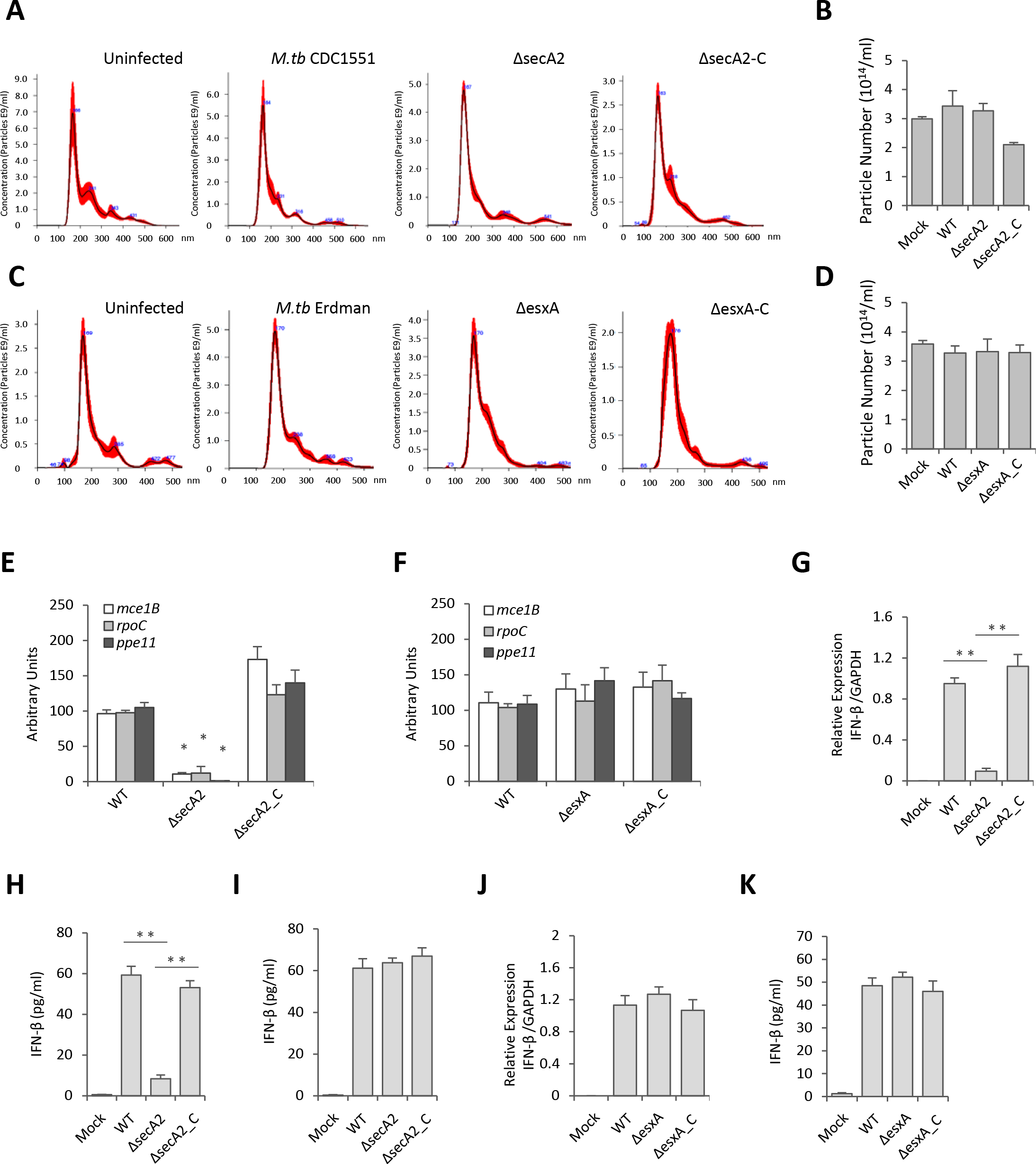
EVs-induced Type I Production in Macrophages Requires the *M.tb* SecA2 Secretion System. **(A)** Nanosight analysis for EVs isolated from BMMs infected with WT, Δ*secA2* or *secA2* complementary (ΔsecA2-C) *M.tb* CDC 1551 strains. **(B)** Yield of purified EVs from BMMs infected with various *M.tb* CDC 1551 strains based on Nanosight analysis. **(C)** and **(D)** As above except using EVs from cells infected with WT, Δ*esxA* or *esxA* complementary (ΔesxA-C) *M.tb* Erdman strains. qRT-PCR analysis for *M.tb* RNA in EVs from BMMs infected with various *M.tb* CDC 1551 strains **(E)** or Erdman strains **(F)**. ND, not detected. qRT-PCR analysis for IFN-β mRNA **(G)** or IFN-β protein **(H)** levels in WT BMMs treated with EVs for 4 hr and 24 hr respectively. The EVs were isolated from BMMs that were infected with the different *M.tb* CDC 1551 strains. **(I)** ELISA analysis for IFN-β secreted by BMMs treated with EVs plus liposomes containing *M.tb* RNA. qRT-PCR analysis for IFN-β mRNA **(J)** or protein **(K)** levels in WT BMMs treated with EVs for 4 and 24 hr respectively. The EVs were isolated from BMMs that were infected with the different *M.tb* Erdman strains. Data shown in (B), (D) and (E - K) are the mean ± SD (n = 3 per group) and representative of three independent experiments. * p < 0.05, ** p < 0.01 and *** p < 0.001 by Student’s t-test (two tailed).

### EVs Released by *M.tb*-infected BMMs Restrict *M.tb* Replication in Host Cells by Activating the *M.tb* RNA/RIG-I/MAVS Signaling Pathway

To test the contribution of EVs in the control of *M.tb* infection, we measured *M.tb* CFU in BMMs pre-treated with EVs from uninfected or *M.tb*-infected macrophages. EVs from *M.tb*-infected macrophages had no significant effect on *M.tb* replication in BMMs in the absence of IFN-γ (Fig. 3A). In contrast, *M.tb* numbers were significantly lower in BMMs pre-treated with EVs released by *M.tb*-infected macrophages in combination with IFN-γ (Fig. 3B). A key survival strategy for *M.tb* is its capacity to inhibit phagosome maturation within infected macrophages (Philips and Ernst, 2012). To begin evaluating the *M.tb* compartment post EV treatment, we stained for the late endosome/lysosome marker Lamp-1. We found elevated colocalization of *M.tb* with Lamp-1 when infected macrophages were pre-treated with IFN-ɣ plus EVs from *M.tb*-infected macrophages relative to IFN-ɣ plus EVs from uninfected macrophages (Fig. 3C and 3D). The rate of Lamp-1 colocalization was comparable in *M.tb*-infected BMMs that were untreated or pre-treated with EVs from uninfected macrophages. To exam whether the host cytosolic RNA sensing pathway plays a role in the EVs-induced phagosome maturation, *M.tb* trafficking was assessed in *Mavs* ^−/−^ BMMs pre-treated with EVs. As shown in Fig. 3E and 3F, the elevated *M.tb* colocalization with Lamp-1 in BMMs pre-treated with IFN-ɣ and EVs from *M.tb*-infected macrophages was absent when using *Mavs* ^−/−^ BMMs. Diminished Lamp-1 colocalization was also seen in RIG-I-knockdown *M.tb*-infected BMMs following pre-treatment with IFN-ɣ and EVs from *M.tb*-infected macrophages (Fig. 3G-3J). Consistent with the Lamp-1 colocalization, *M.tb* burden in *Mavs* ^−/−^ BMMs and RIG-I siRNA-treated BMMs was higher relative to control BMMs when cells were pre-pretreated with IFN-ɣ and EVs from *M.tb*-infected macrophages (Fig. 3B and 3K-3M). However, even in the absence of the RNA signaling pathway, EVs from *M.tb*-infected BMMs reduce *M.tb* burden within infected cells relative to untreated BMMs, indicating that EVs have additional effects on host cells which impacts bacterial survival and/or replication.

**Figure 3.**
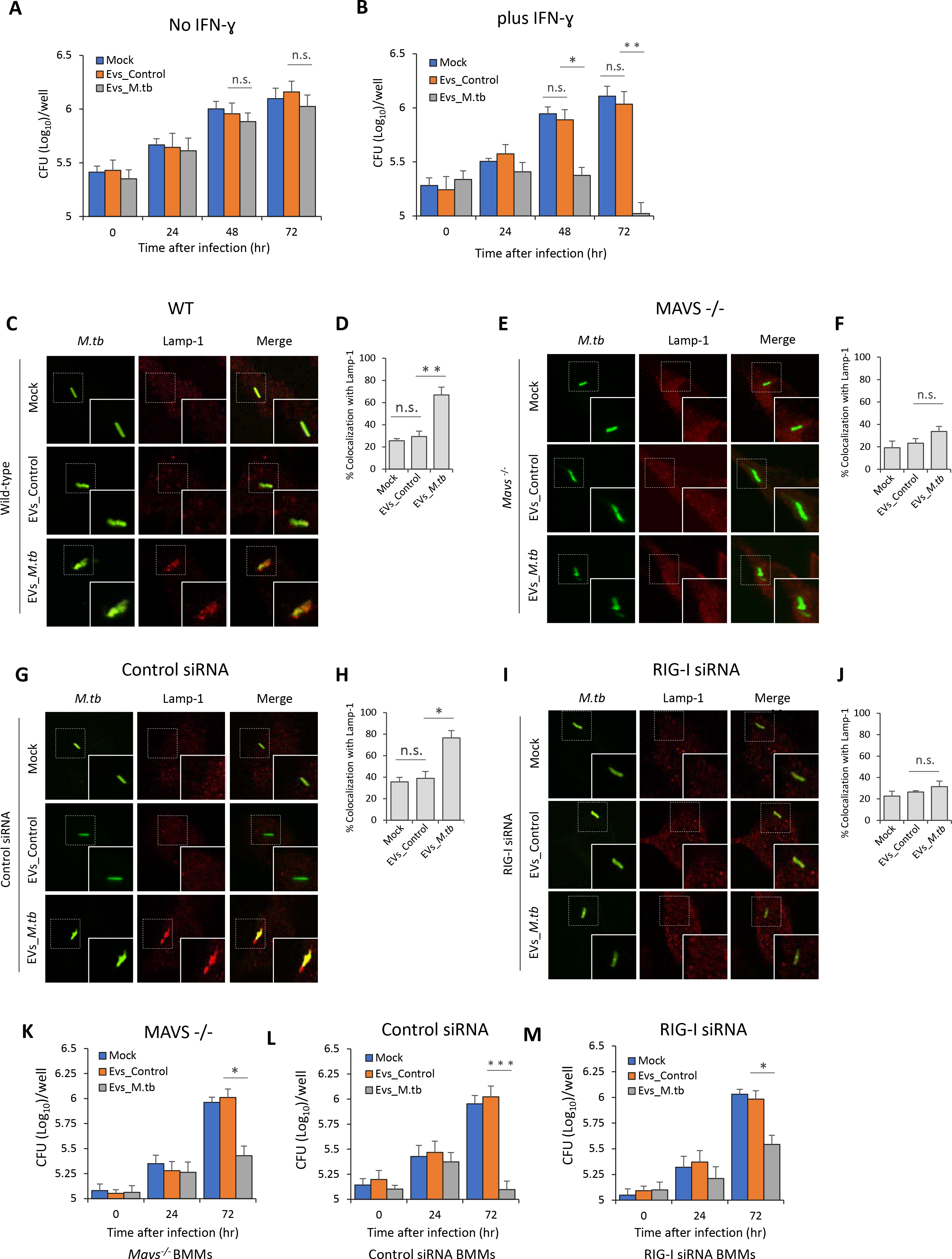
EVs Released by *M.tb*-infected Macrophages Restrict *M.tb* Replication in Host cells by Activating the *M.tb* RNA/RIG-I/MAVS Signaling Pathway. *M.tb* CFU in WT mouse BMMs pre-treated with EVs minus **(A)** or plus **(B)** co-treatment with IFN-ɣ. BMMs were treated with EVs from uninfected (EVs_Control) or *M.tb*-infected (EVs_*M.tb*) cells for 0, 24, 48 and 72 hr following a 1 hr *M.tb* infection. Mock, no EV treatment. Immnofluorescence microscopy analysis for colocalization of *M.tb* (GFP) with lysosome marker Lamp-1 in WT **(C)** or *Mavs^−/−^* **(E)** mouse BMMs. Cells were pre-treated for 5 hr with EVs from uninfected or *M.tb*-infected macrophages plus IFN-γ, and then infected with GFP-expressing *M.tb* for 24 hr prior to immunostaining. Immunofluorescence microscopy as described above except using control siRNA-treated **(G)** or RIG-I siRNA-treated **(I)** BMMs. **(D)** and **(F)** quantitative analysis of *M.tb* colocalization with Lamp-1 in WT and *Mavs^−/−^* mouse BMMs respectively. **(H)** and **(J)** quantitative analysis of *M.tb* colocalization with Lamp-1 in control siRNA-treated or RIG-I siRNA-treated mouse BMMs respectively. *M.tb* CFU in infected *Mavs*^*−/−*^ **(K)**, control siRNA-treated **(L)** or RIG-I siRNA-treated **(M)** mouse BMMs pre-treated with IFN-ɣ plus EVs from uninfected (EVs_Control) or *M.tb*-infected (EVs_*M.tb*) BMMs. CFU was determined immediately after the 1 hr infection or 24 and 72 hr post-infection. Data shown are the mean ± SD (n = 3 per group) and representative of at least three independent experiments. n.s., not significant; * p < 0.05, ** p < 0.01 and *** p < 0.001 by two-tailed Student’s t-test.

To investigate whether the *M.tb* RNA in EVs is required for the RIG-I/MAVS-dependent *M.tb*-killing pathway, WT BMMs were pre-treated with EVs isolated from macrophages that were infected with WT, Δ*secA2* or *secA2* complementary strains. In contrast to EVs from WT or *secA2* complementary strain-infected macrophages, EVs from Δ*secA2 M.tb*-infected macrophages failed to promote phagosome maturation and suppress *M.tb* replication in BMMs (Fig. 4A, 4B and 4E). This deficiency of EVs from Δ*secA2 M.tb*-infected macrophages was rescued by adding liposome-encapsulated RNA, with the RNA isolated from EVs released from M.tb-infected BMMs. Adding this RNA resulted in increased phagosome maturation and decreased *M.tb* survival (Fig. 4C, 4D and 4F) indicating that EVs-associated *M.tb* RNA was driving the anti-mycobacterial response in recipient cells.

**Figure 4.**
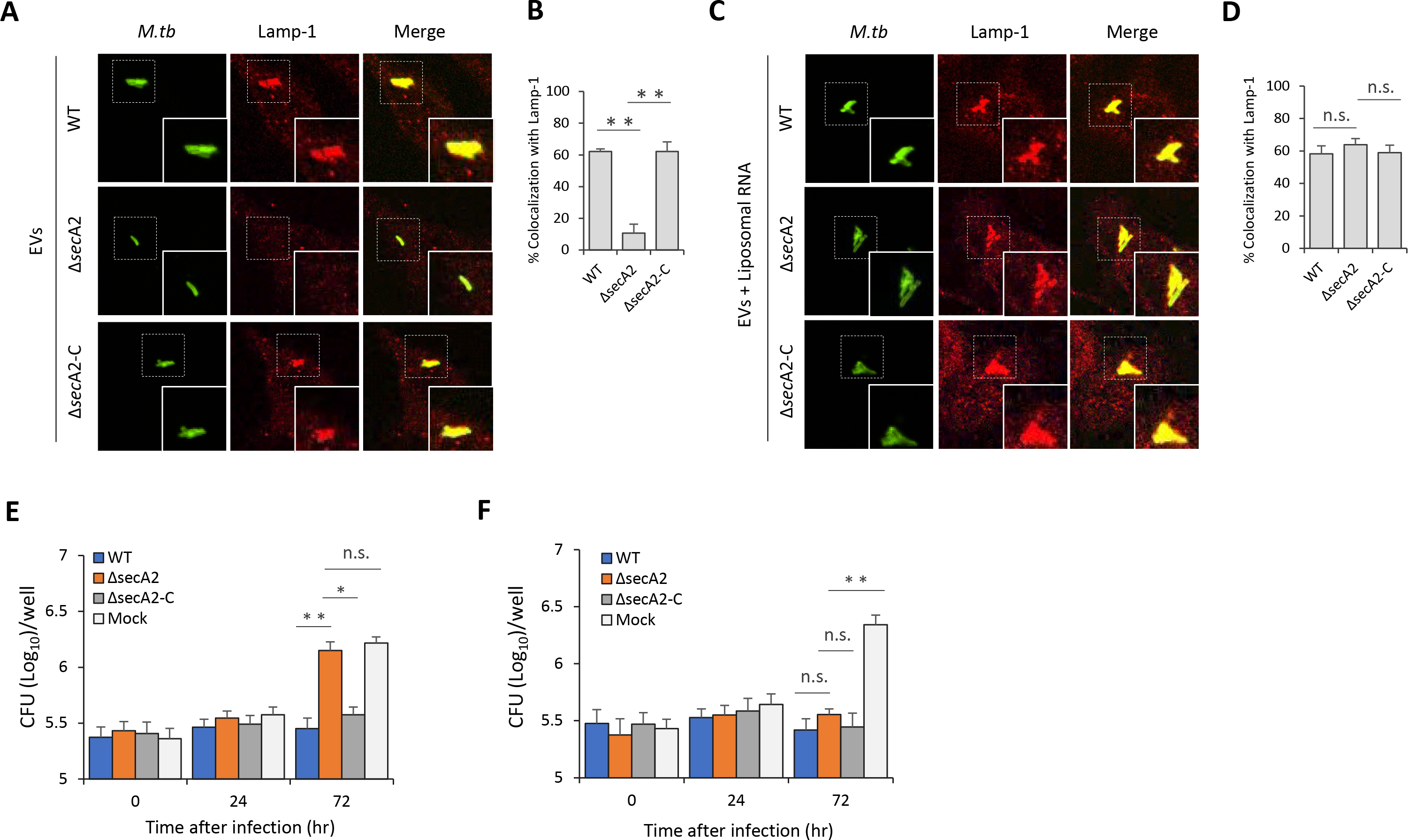
EVs-stimulated *M.tb* Phagolysosome Maturation in BMMs is dependent on the Mycobacterial SecA2 Secretion System. **(A)** Immunofluorescence microscopy analysis for colocalization of *M.tb* with Lamp-1 in WT BMMs pre-treated with EVs from macrophages infected with WT, Δ*secA2* or *secA2* complementary (ΔsecA2-C) *M.tb* CDC 1551 strains. The cells were pre-treated with EVs supplemented with recombinant mouse IFN-γ for 5 hr and subsequently infected for 24 hr with GFP-expressing *M.tb*. **(B)** quantitative analysis for the colocalization of *M.tb* with Lamp-1. **(C)** and **(D)** as described above but liposomes containing RNA isolated from EVs released from BMMs infected with WT *M.tb* were also added to cells 5 hr prior to infection. **(E)** *M.tb* CFU in WT BMMs pre-pretreated with recombinant mouse IFN-γ and EVs from macrophages infected with WT, Δ*secA2* or *secA2* complementary (ΔsecA2-C) *M.tb* CDC 1551 strains. **(F)** As above but again liposome-encapsulated RNA isolated from EVs released from BMMs infected with WT *M.tb* was included during EV and IFN-γ pre-treatment. Data shown in (B), (D), (E) and (F) are the mean ± SD (n = 3 per group) and representative of at least three independent experiments. n.s., not significant; * p < 0.05 and ** p < 0.01 by two-tailed Student’s t-test.

### EVs Released by *M.tb*-infected BMMs Activate LC3-associated Phagocytosis Pathway in BMMs during *M.tb* Infection

Autophagy plays a key role in the clearance of intracellular pathogens. Recently, it was found that ubiquitin (Ub)-mediated autophagy contributes to the control of *Mycobacterium bovis* BCG and *M.tb* infection in host cells through a TBK1-regulated pathway (Pilli et al., 2012; Watson et al., 2012). The MAVS-dependent activation of TBK1 by EVs from *M.tb*-infected macrophages suggest that these EVs may regulate this autophagy pathway. To test this hypothesis, the colocalization of *M.tb* with the autophagosome biomarker LC3 and Ub was investigated by immunofluorescence microscopy. As shown in Fig. 5, pre-treatment of control siRNA-treated (Fig. 5A and 5B) or WT (Fig. 5E and 5F) BMMs with EVs from *M.tb*-infected cells plus IFN-ɣ significantly increased colocalization of *M.tb* with LC3 compared to the untreated BMMs or BMMs treated with EVs from uninfected macrophages. This increased phagosome maturation was not seen in either RIG-I-knockdown (Fig. 5C and 5D) or *Mavs* ^−/−^ BMMs (Fig. 5G and 5H). In contrast, neither EVs from uninfected or from *M.tb*-infected BMMs promoted the trafficking of *M.tb* into Ub-positive vesicles in BMMs (Fig. S2A). Furthermore, a knockdown of TBK1 had no significant effect on the colocalization of *M.tb* with LC3-positive vesicles in WT BMMs treated with IFN-ɣ plus EVs (Fig. S2B and S2C), suggesting an alternative LC3-dependent autophagic pathway.

**Figure 5.**
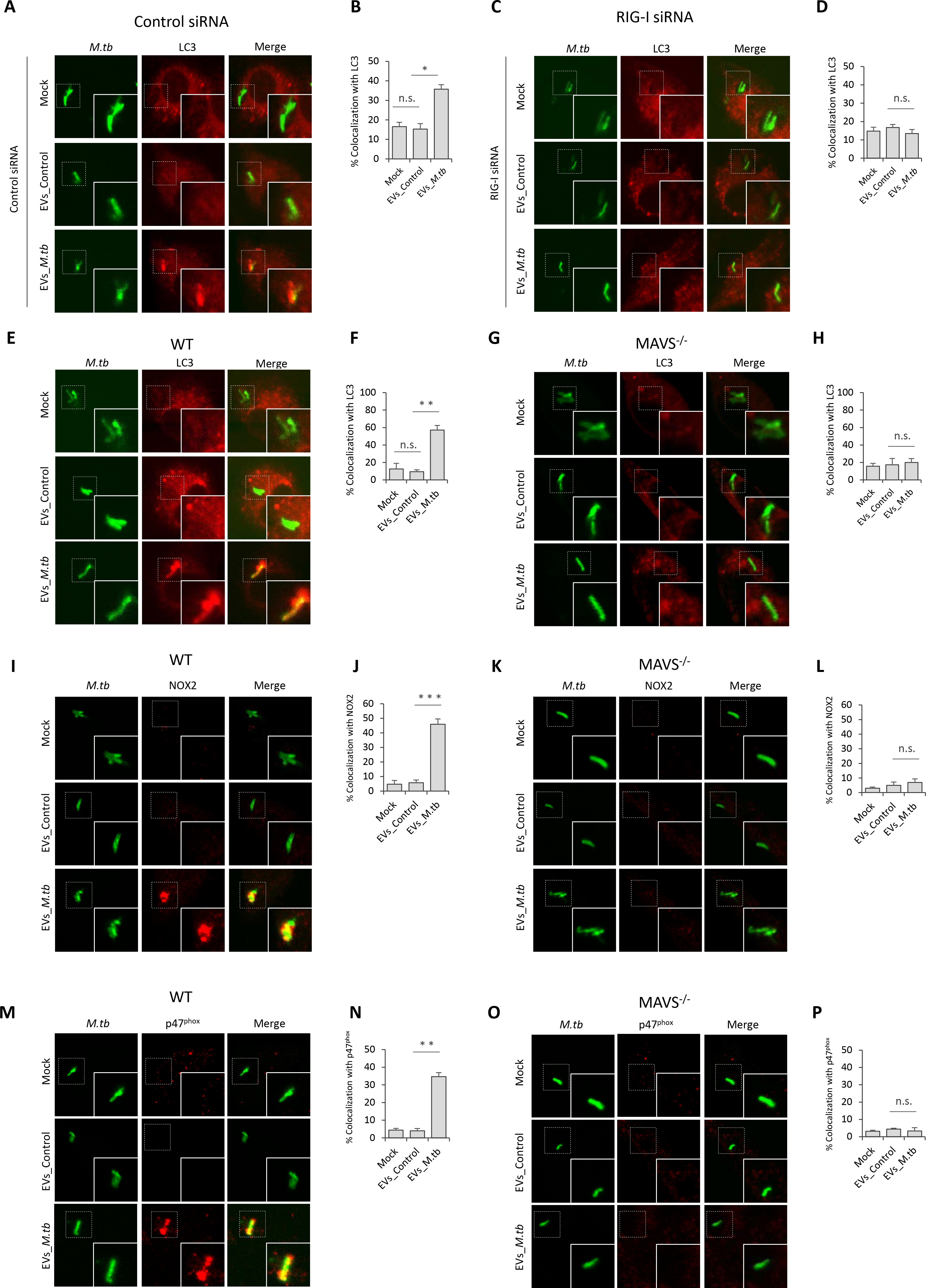
EVs Released by *M.tb*-infected Macrophages Activate LC3-associated Phagosome Maturation via a RIG-I/MAVS-dependent pathway in BMMs. Immunofluorescence microscopy analysis for colocalization of *M.tb* with autophagosome marker LC3 in control siRNA **(A)** or RIG-I siRNA **(C)** treated BMMs that were untreated or pre-treated for 5 hr with recombinant mouse IFN-γ and EVs from uninfected (EVs_Control) or *M.tb*-infected (EVs_*M.tb*) macrophages followed by a 24 hr infection with GFP-expressing *M.tb*. Mock, untreated. **(B)** and **(D)** quantitative analysis of this *M.tb* colocalization with LC3. Immunofluorescence microscopy analysis for colocalization of *M.tb* with markers LC3 **(E)** NOX2 **(I)** or p47^phox^ **(M)** in WT BMMs that were left untreated or pre-treated for 5 hr with recombinant mouse IFN-γ and EVs from uninfected (EVs_Control) or *M.tb*-infected (EVs_*M.tb*) macrophages, followed by a 24 hr infection with GFP-expressing *M.tb*. Mock, untreated. Immunofluorescence microscopy analysis for colocalization of *M.tb* with LC3 **(G)**, NOX2 **(K)** and p47^phox^ **(O)** as described above but using *Mavs* ^−/−^ BMMs. Quantitative analysis for colocalization of *M.tb* with LC3 **(F)**, NOX2 **(J)** and p47^phox^ **(N)** in WT BMMs. Quantitative analysis for colocalization of *M.tb* with LC3 **(H)**, NOX2 **(L)** and p47^phox^ **(P)** in *Mavs* ^−/−^ BMMs. Quantitative data are the mean ± SD and representative of at least three independent experiments. n.s., not significant; * p < 0.05, ** p < 0.01 and *** p < 0.001 by two-tailed Student’s t-test.

LC3-associated phagocytosis (LAP), a Ub-independent process, was recently uncovered in the host defense against bacterial infection (Martinez et al., 2015). After engagement/activation of the PRR by the bacterial PAMP, NOX2 NADPH oxidase complex was recruited to the phagosomal membrane to stimulate the production of reactive oxygen species (ROS), which subsequently promoted recruitment of LC3 to the phagosome, facilitating phagosome-lysosome fusion (Huang et al., 2009). The NOX2 NADPH oxidase constitutes a membrane-bound subunit (NOX2/gp91*^phox^*, and p22*^phox^*) and three cytosolic components p67*^phox^*, p47*^phox^*, and p40*^phox^*. To test whether LAP is involved in EVs-triggered phagosome-lysosome fusion in *M.tb*-infected BMMs, we analyzed the colocalization of NOX2 and p47*^phox^* with *M.tb* in BMMs. Similar to LC3, treatment of BMMs with EVs from *M.tb*-infected macrophages significantly increased colocalization of *M.tb* with NOX2 (Fig. 5I and 5J) and p47*^phox^* (Fig. 5M and 5N). This effect of EVs relies on the host RNA sensing pathway as this increased colocalization of *M.tb* with NOX2 (Fig. 5K and 5L) and p47*^phox^* (Fig. 5O and 5P) was not observed in *Mavs* ^−/−^ BMMs.

### EVs Released by *M.tb*-infected BMMs Synergistically Attenuate *M.tb* Survival in BMMs When Combined with Moxifloxacin

The ability of EVs from *M.tb*-infected macrophages to inhibit *M.tb* survival in host cells suggest these vesicles may have some potential in anti-TB therapy. To test this hypothesis, WT mouse BMMs were first infected with wild type *M.tb*, and 24 hr post-infection, cells were treated with a combined regimen consisting of EVs from *M.tb*-infected macrophages and moxifloxacin, a key antibiotic against MDR-TB. As shown in Fig. 6A, an EVs-moxifloxacin combination significantly increased *M.tb* trafficking to a Lamp-1 positive compartment compared to EVs or moxifloxacin alone. In contrast, in *Mavs* ^*−/−*^ BMMs, EVs from *M.tb*-infected macrophages failed to enhance the effect of moxifloxacin as a similar number of *M.tb* colocalized with Lamp-1 in cells treated with moxifloxacin alone compared to the EVs-moxifloxacin combination (Fig. 6B). Moreover, moxifloxacin combined with EVs from *M.tb*-infected macrophages also resulted in increased trafficking of *M.tb* to LC3 (Fig. 6C), NOX2 (Fig. 6E) and p47^*phox*^ (Fig. 6G) positive compartments when compared to moxifloxacin or EVs-treatment alone. Additional studies indicated that the effect of these EVs on *M.tb* trafficking to LC3 (Fig. 6D), NOX2 (Fig. 6F) and p47*^phox^* (Fig. 6H) positive compartments was MAVS dependent. An effect on *M.tb* survival was also observed within infected BMMs as EVs from *M.tb*-infected macrophages significantly enhanced the efficacy of moxifloxacin (Fig. 6I). Consistent with the Lamp-1 colocalization results, no difference in bacterial load was detected in *Mavs* ^*−/−*^ BMMs between moxifloxacin alone and when the drug was combined with EVs from *M.tb*-infected macrophages (Fig. 6J). Similar to the EV pretreatment studies, there was a similar level of colocalization between Ub and *M.tb* in untreated BMMs compared to post-exposure treatments with EVs (Fig. S3A), and no TBK1 involvement was apparent in the delivery of *M.tb* to a LC3 positive compartment (Fig. S3B).

**Figure 6.**
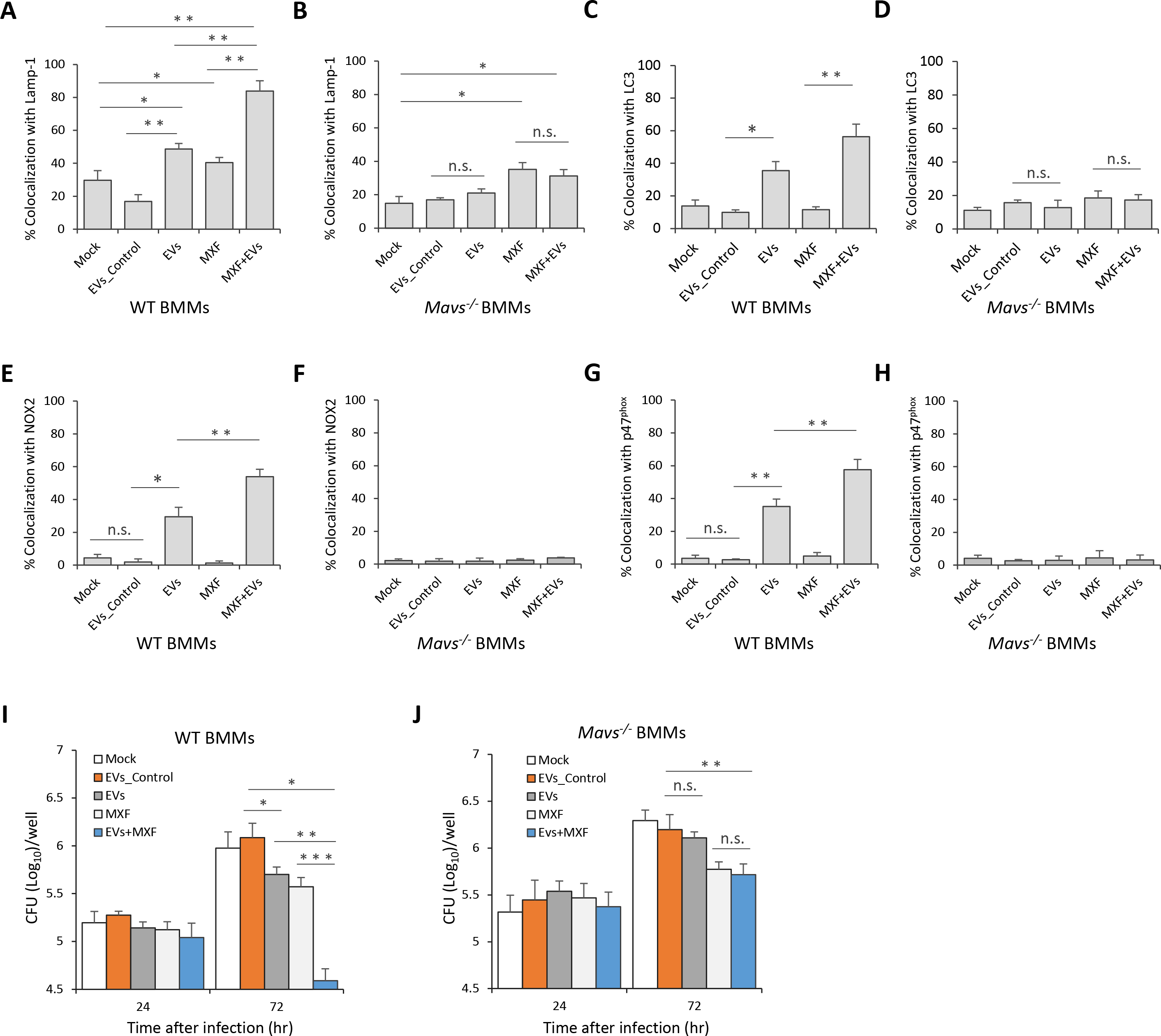
EVs Released by *M.tb*-infected BMMs Attenuate *M.tb* Survival in Macrophages When Combined with Moxifloxacin. Quantitative analysis of immunofluorescence microscopy images for colocalization of *M.tb* with Lamp-1 **(A)**, LC3 **(C)**, NOX2 **(E)** and p47^phox^ **(G)** in WT BMMs infected with *M.tb* for 24 hr and then treated for an additional 24 hr with moxifloxacin (MFX) and/or EVs from *M.tb*-infected BMMs (EVs+MFX). Mock, no EVs or moxifloxacin treatment; EVs_Control, EVs from uninfected BMMs. As described above except *Mavs* ^−/−^ BMMs were used and colocalization of *M.tb* with Lamp-1 **(B)**, LC3 **(D)**, NOX2 **(F)** and p47^phox^ **(H)** was quantified. *M.tb* CFU analysis in WT **(I)** and *Mavs* ^−/−^ **(J)** BMMs infected with *M.tb* for 24 hr followed by treatment with EVs, moxifloxacin or combination for 24 and 72 hr. Data shown are representative of three independent experiments. The results are the mean ± SD (n = 3 per group). n.s., not significant; * p < 0.05, ** p < 0.01 and *** p < 0.001 by Student’s t-test (two tailed).

### EVs Released by *M.tb*-infected BMMs Significantly Decreased *M.tb* Survival in Mice When Combined with Moxifloxacin

The decreased bacterial numbers observed in WT BMMs after treatment with moxifloxacin plus EVs from *M.tb*-infected macrophages suggest that host cell-derived EVs might be effective immunotherapy in combination with anti-TB drugs. To test this hypothesis, WT C57BL/6 mice were low-dose aerosol-infected with *M.tb*, which was followed 3 weeks later with a two-week treatment with moxifloxacin and a single dose EV treatment given 4 weeks post-infection (Fig. 7A). As seen in Fig. 7B, mice treated with moxifloxacin or EVs from *M.tb*-infected macrophages, or combination therapy had smaller granuloma-like lesions in the lung when compared to untreated mice or those receiving EVs from uninfected macrophages. Consistent with histopathological results, these groups of mice had significantly lower mycobacterial burden in the lung and spleen (<u>Fig. 7C</u>). Interestingly, moxifloxacin-EVs combined treatment was more effective than moxifloxacin or EVs alone (Fig. 7B and 7C). To determine whether EVs-based immunotherapy is dependent on MAVS, we performed the combination treatment using *M.tb*-infected *Mavs* ^*−/−*^ mice. Consistent with the in vitro results using BMMs, EVs from *M.tb*-infected macrophages failed to boost moxifloxacin-based chemotherapy in *M.tb*-infected *Mavs* ^*−/−*^ mice. No significant histopathological difference was seen between the lungs of the different *M.tb*-infected groups in *Mavs* ^*−/−*^ mice (Fig. 7D) and a similar *M.tb* count was seen in the lung and spleen of *Mavs* ^*−/−*^ mice receiving only moxifloxacin compared to mice treated with the combination of moxifloxacin and EVs (<u>Fig. 7E</u>). Cytokine levels were also affected by EVs as higher levels of IFN-β was found in the serum of *M.tb*-infected mice following treatment with EVs from *M.tb*-infected macrophages (Fig. 7F). This EV-stimulated IFN-β production in mice was dependent on MAVS (Fig. 7G). Finally, EVs from *M.tb*-infected macrophages also induced increased levels of TNF-α and IL-1β production in *M.tb*-infected mice via a MAVS-independent pathway but had no effect on IFN-γ production during the infection period (Fig. 7F and 7G).

**Figure 7.**
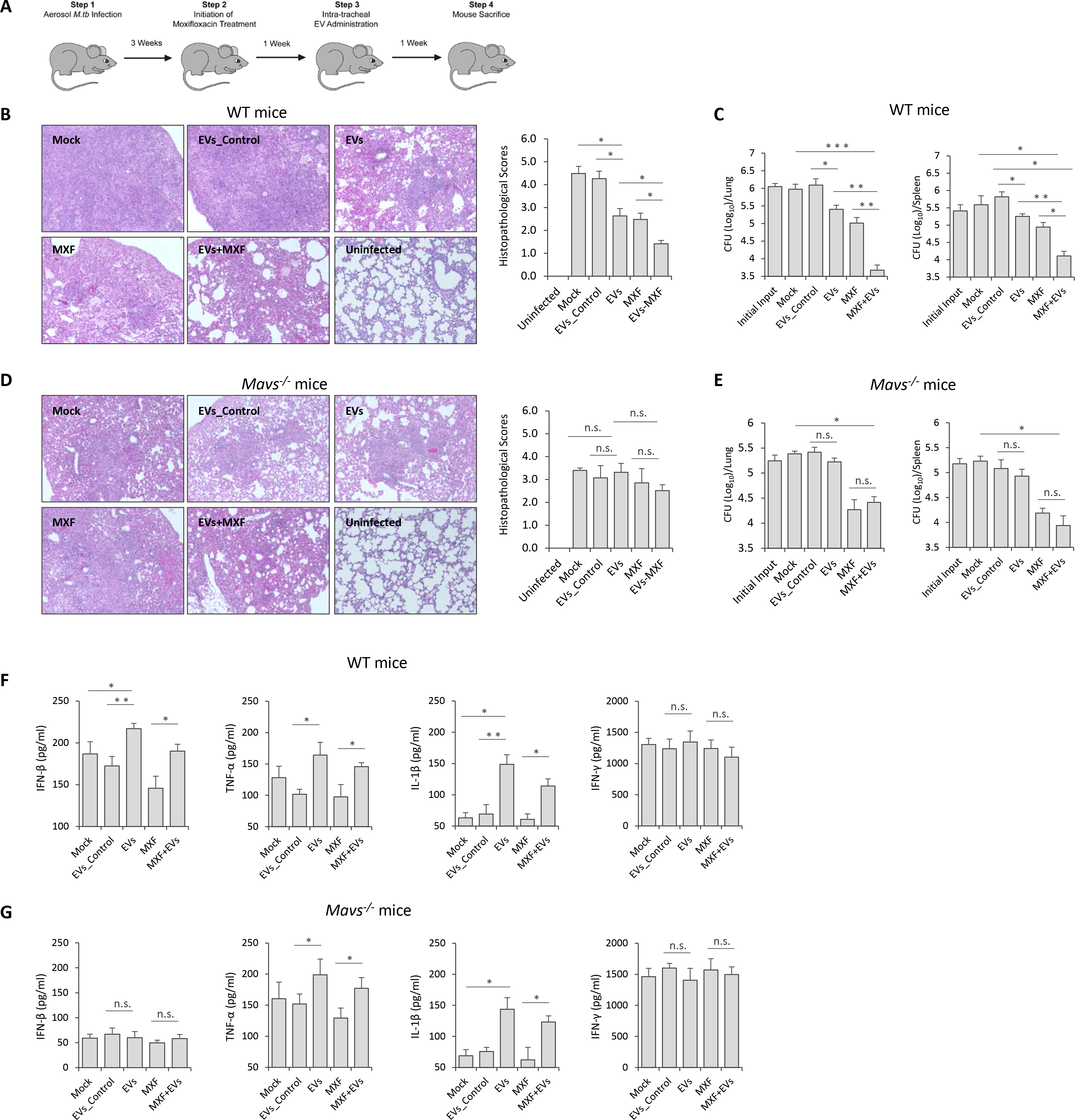
EVs Released by *M.tb*-infected Macrophages Significantly Decreased *M.tb* Survival in Mice When Combined with Moxifloxacin. **(A)** Schematic for EVs-based adjunctive immunotherapy and moxifloxacin-based chemotherapy in *M.tb*-infected mice. Representative histopathological analysis for lung sections of WT **(B)** and *Mavs* ^−/−^ **(D)** mice that were infected with *M.tb* and subsequent left untreated (Mock) or treated with EVs from uninfected BMMs (EVs_Control), EVs from *M.tb*-infected BMMs (EVs), moxifloxacin (MXF), or combination of EVs and MXF (EVs+MXF). *M.tb* CFU in the lung and spleen of WT **(C)** or *Mavs* ^−/−^ **(E)** mice treated with EVs, MXF or combination of both. ELISA analysis for, IFN-β, TNF-α, IL-1β and IFN-γ protein level in serum isolate from *M.tb*-infected WT **(F)** or *Mavs* ^−/−^ **(G)** mice treated with EVs, MXF or a combination of both. Data shown is representative of two independent experiments. The results in (B) – (G) are the mean ± SD (n = 4 per group). n.s., not significant; * p < 0.05, ** p < 0.01 and *** p < 0.001 by Student’s t-test (two tailed).

## Discussion

Cell-to-cell communication plays a critical role in host defense against microbial infections. For intracellular pathogens, communication between infected host cells and cells of the immune system is mediated through cell-cell contact or release of soluble factors by the infected cell including cytokines, chemokines and various inflammatory mediators. Recently, EVs are recognized as key players in intercellular communication and may transfer pathogen-derived nucleic acids and proteins to bystander cells. However, there remains limited information on how these EVs modulate the host response to infection (Schorey et al., 2015). Previous studies from our laboratory and others have begun to characterize the role of EVs in intercellular communication during an *M.tb* infection using both infected macrophages and mouse infection models. We found that EVs from *M.tb*-infected macrophages induce the production of multiple cytokines including TNF-α in recipient cells through a MyD88-dependent pathway (Singh et al., 2012). In the present study, we found that EVs containing *M.tb* RNA may deliver bacterial nucleic acids into uninfected cells, leading to the activation of the RIG-I/MAVS-dependent RNA sensing pathway. Together these data suggest that various *M.tb* PAMPs or host signal molecules are carried in EVs from *M.tb*-infected macrophages, and these molecules dictate the effect of EVs on recipient cells. However, the lack of a suitable animal model that is impaired or deficient in EV biogenesis has hampered the *in vivo* studies to address the positive or negative effect of EVs on infection.

Our study indicates that *M.tb* RNA released during a macrophage infection requires expression of the mycobacterial SecA2 protein. Unlike SecA1, SecA2 is dispensable for growth and exports only a limited number of proteins. These SecA2-dependent secreted proteins are involved in bacterial pathogenesis and cellular responses to environmental stress (Feltcher and Braunstein, 2012). The SecA2 protein has been identified in all mycobacterial strains and some Gram-positive bacteria including *Bacillus*, *Clostridium*, *Corynbacteria*, *Listeria*, *Staphylococcus*, and *Streptococcus* species (Green and Mecsas, 2016). In *L. monocytogenes*, the deficiency of the SecA2 protein significantly decreases bacterial RNA release during bacterial culture (Abdullah et al., 2012). In *M.tb*, we also found that the SecA2 protein is critical for mycobacterial RNA release during growth in culture media (data not shown). These results suggest that the secretion of bacterial RNA into the extracellular environment might exist as a ubiquitous pathway for bacteria expressing a SecA2-secretion system. Moreover, although we only evaluated the transfer of bacterial RNA to EVs during the course of an *M.tb* infection, it is possible that the intercellular transfer of bacterial RNA via host cell-derived EVs is also observed for other pathogens that express a SecA2 expression system.

EVs from *M.tb*-infected macrophages promoted phagosome maturation in *M.tb*-infected macrophages when used as pre-treatment agent or after an *M.tb* infection, leading to reduced mycobacterial replication. Our results also found that EVs from *M.tb*-infected macrophages trigger *M.tb*-containing phagosome maturation through a LC3-associated pathway (Mitchell and Isberg, 2017). LAP represents an alternative autophagy-dependent antimicrobial pathway in host cells, in which LC3-modified vesicles fuse with lysosomes, promoting microbial degradation (Mitchell and Isberg, 2017). Unlike classical autophagy, LAP is a Ub-independent process and only utilizes a subset of autophagy machinery components for the modification of microbe-containing vesicles by the LC3-conjugation system (Lam et al., 2013; Hubber et al., 2017). As an established intracellular bacterial pathogen, *M.tb* has evolved an inhibitory mechanism for evading LAP through release of CpsA, a LytR-CpsA-Psr (LCP) domain-containing protein that may interfere with the recruitment of NOX2 NADPH oxidase to *M.tb*-containing phagosomes (Köster et al., 2017). Interestingly, EVs-mediated LC3 conjugation of *M.tb*-containing phagosomes requires the host RIG-I/MAVS cytosolic RNA sensing pathway. Our study highlights a previously undefined role for the host RNA sensing pathways in noncanonical LC3-associated phagosome maturation in host cells during the course of an *M.tb* infection.

Drug-resistant TB is becoming a major threat in the global TB control (WHO Report, 2017). Globally in 2016, MDR/RR-TB was diagnosed in an estimated 4.1% of new cases and about 19% of previously treated cases. Among these, approximately 6.2% of cases was XDR-TB. An estimated treatment success rate for MDR/RR-TB and XDR-TB was 54% and 30%, respectively. Treatment for MDR/RR-TB and XDR-TB requires a longer therapeutic duration with less effective, more expensive and higher toxicity drugs, leading to a high rate of treatment failure and mortality. To stop the global spread of MDR/RR-TB and XDR-TB, new anti-TB drugs or combined regimens are urgently needed. Recently, a combined therapeutic strategy consisting of an adjunct immunotherapy and antimycobacterial drugs has been proposed and investigated (Uhlinf et al., 2012). The agents most commonly used in TB immunotherapy include various immune mediators such as all-trans retinoic acid which is known to deplete myeloid-derived suppressor cells as well as increase expression of CD1d on antigen presenting cells. When all-trans retinoic acid in combination with the CD1b ligand alpha-galactosylceramide was administered to mice along with antibiotics there was significant improvement in bacterial clearance and lower relapse rates than when treated with isoniazid, rifampicin, and pyrazinamide alone (Mourik et al., 2016). Other immune-based therapy have targeted various pathways that are responsible for driving a host inflammatory response following a mycobacterial infection with the goal of promoting the right balance between too little and too much inflammation (Kiran, et al., 2016). Such host targets include inhibiting PGE2 production which is associated with increased IL-10 production. Recent work has also focused on ways to stimulate angiogenesis promoting increased blood supply which will allow for more efficient drug penetration and increase the access of host immune cells to the granuloma (Dartois, 2014).

In the present study, we investigated an alternative approach that consist of a EVs-based immunotherapy combined with a mycobacterial antibiotic. Unlike the agents investigated previously, the EVs derived from *M.tb*-infected host cells will have a more limited target cell population as prior studies indicate a predisposition for EV uptake by macrophage and DCs (Bhatnagar et al., 2007). Targeting mycobacterial PAMPs and antigens into *M.tb*-infected macrophages or uninfected host cells may trigger antimycobacterial pathways such as LAP as well provide *M.tb* antigens for activation of an acquired immune response (Bhatnagar et al., 2007; Köster et al., 2017). We found that EVs containing *M.tb* PAMPs such as mycobacterial RNA are able to elicit an effective antimycobacterial response in macrophages. This suggested that EVs may promote clearance *in vivo* and provide additional benefit to antibiotic treatment. Indeed we observed both decreased bacterial load and limited lung pathology in *M.tb*-infected mice treated with EVs and moxifloxacin compared to either alone. Future studies to identify the EVs surface molecules that are responsible for EVs-cell recognition and contact, could potentially guide development of artificial particles carrying anti-TB immunotherapeutic agents that are targeted to the appropriate macrophage population potentially increasing its efficacy. Previous studies have supported this concept that EVs, which are targeted to specific cell populations, can have significantly higher therapeutic activity. For example, exosomes isolated from HEK293 cells were pre-loaded with synthetic let-7a miRNA, a tumor suppressor. When these exosome’s expressed the transmembrane domain of platelet-derived growth factor receptor fused to the GE11 peptide, they were specifically targeted to xenograft breast cancer cells via a GE11-EGFR interaction resulting in reduced tumor growth in *RAG2* ^−/−^ mice (Ohno et al., 2013). Our study extends the potential application of EVs as immunotherapeutic agents, especially as an adjunctive therapy for currently drug resistant infections caused by intracellular pathogens such as *M.tb*.

In summary, we found that the presence of *M.tb* RNA in EVs released from infected macrophages is dependent on the bacteria SecA2 secretion system. Further, these EVs carrying *M.tb* RNA can activate the host RIG-I/MAVS/TBK1/IRF3 RNA sensing signaling pathway in recipient macrophages, leading to the production of type I IFNs. Additionally, a RIG-I/MAVS-dependent phagosome maturation is induced by EVs from *M.tb*-infected macrophages, resulting in an increased trafficking of *M.tb* into LC3 and Lamp-1 positive vesicles and increased bacterial killing. Finally, we found that EVs can synergize with TB antibiotics to promote bacterial clearance and limit lung pathology suggesting a novel immuotherapeutic approach to treat drug-resistant *M.tb*.

## Experimental Procedures

### Mice

Wild type C57BL/6 and *MyD88* ^−/−^ mice have been described previously (Bhatnagar et al., 2007). *Mavs*^−/−^ mice on a C57BL/6 background were obtained as a generous gift from Dr. Stanley Perlmanb (University of Iowa, USA) (Suthar et al., 2012). All mice were housed at the institutional animal facility under specific pathogen-free conditions during the experiment. The University of Notre Dame is accredited through the Animal Welfare Assurance (#A3093-01). All animal experiments were approved by the Institutional Animal Care and Use Committees (IACUCs) of University of Notre Dame.

### Bacterial Strains

All *M.tb* strains were grown in MiddleBrook 7H9 broth (Cat.271310, BD) supplemented with 10% (v/v) Middlebrook oleic acid-albumin-dextrose-catalase (OADC) (Cat. 211886, BD) and 0.2% glycerol until mid-exponential phase, and washed with complete medium for macrophages or ddH2O plus 0.05% Tween-80 when required.

### Cell Culture

Bone Marrow-derived Macrophages (BMMs) were isolated from wild type C57BL/6, *Mavs*^−/−^ or *MyD88* ^−/−^ mice (female, 6-8 weeks) as described previously (Roach and Schorey, 2002), and cells were grown in DMEM supplemented with 10% (v/v) heat-inactivated FBS, 20% L929 cell-conditional medium as a source of macrophage colony-stimulating factor and 100 U/ml penicillin and 100 U/ml streptomycin (SV30010, HyClone) at 37°C and 5% CO_2_.

### siRNA Transfection

Mouse BMMs (3 × 10^5^ cells/well) were transfected with AllStars Negative Control siRNA (Cat.1027280, Qiagen), RIG-I (5’-GAAGCGUCUUCUAAUAAUU-3’), MAVS (5’-GAUCAAGUGACUCGAGUUU-3’ and 5’-GGACCAAAUAGCAGUAUCA-3’) and TBK1 (SMARTpool: ON-TARGETplus Tbk1 siRNA, Dharmacon) siRNA oligos (25 pmol/3 × 10^5^ cells) in 24-well plates using Lipofectamine 2000 (Cat.11668-027, Invitrogen) following the manufacturer’s protocol. The transfected cells were cultured in BMM complete medium for 48 hr before.

### Macrophage-derived EVs Isolation

BMMs were infected with various *M.tb* strains at an MOI of 5 for 4 hr and washed with pre-warm PBS (1x) three times to remove remaining *M.tb*. Infected cells were incubated in DMEM supplemented with 10% EVs-free FBS for an additional 72 hr and exosome-enriched EVs were isolated as described previously (Cheng et al., 2013). Isolated EVs were quantified using BCA protein assay and the NanoSight LM10 (Malvern Panalytical, UK).

### Survival Assay of *M.tb* Strains in BMMs

For EV pretreatment experiments, BMMs were treated with EVs at 10 μg/ml for 5 hr, and subsequently infected with *M.tb* strains at an MOI of 5 for 1 hr at 37°C and 5% CO_2_. The cells were then washed with complete BMMs medium 3 times, and further incubated for another 24, 48 and 72 hr at 37°C and 5% CO_2_. Finally, cells were washed with pre-cold PBS 3X and lysed in 0.05% SDS. A series of dilution of cell lysates in PBS (1x) were added onto 7H11 agar plates (Cat.7244A, Acumedia) supplemented with 10% (v/v) OADC and 0.2% glycerol. Plates were incubated at 37°C for 3-4 weeks until counting. For EVs and moxifloxacin treatment post *M.tb* infection, BMMs were first infected with *M.tb* for 1 hr at 37°C and 5% CO_2_ and then washed with complete BMMs medium 3 times. The cells were incubated for 24 hr at 37°C and 5% CO_2_ before treatment with EVs at 5 μg/ml and moxifloxacin at 1.0 μg/ml for 24 and 72 hr at 37°C and 5% CO_2_ in the presence of recombinant mouse IFN-γ (200 Units, Cat.No.14-8311-63, Invitrogen).

### Transwell Assay

Wild-type BMMs were infected with WT *M.tb* strain at a MOI of 5 for 1 hr at 37°C and 5% CO_2_, and then washed with complete BMMs medium 3 times. Cells were incubated for another 24 hr at 37°C and 5% CO_2_ before transferring into a transwell inserts (pore size, 0.4μm; Cat. 3413, Corning) that were subsequently co-incubated with WT or *Mavs*^−/−^ BMMs pre-seeded in the lower compartment. The IFN-β mRNA level within BMMs in the lower chamber was analyzed by qRT-PCR.

### RNA Purification

To determine mycobacterial RNA in macrophage-derived EVs, total RNA in EVs was isolated using mirVana™ miRNA Isolation Kit (AM1560, Invitrogen). For IFN-β mRNA measurement, BMMs were treated with isolated EVs at 37°C and 5% CO_2_ for various time as required and then total cellular RNA was purified using Qiagen RNeasy Plus Mini Kit (Cat.No. 74136, Qiagen).

### qRT-PCR

RNAs were initially treated with DNase I (Cat.18068015, Invitrogen) following the manufacturer’s introduction. For mycobacterial RNA in EVs, cDNA were synthesized with AMV Reverse Transcriptase (Cat.M0277, NEB) and a mixture of *M.tb* Reverse primers. *mce1B*: Forward, 5’-GAGATCGGCAAGGTCAAAGC-3’; Reverse: 5’-GCGGTCGTGGACTGATACAA-3’, *rpoC*: Forward, 5’-ATGGTGACCGGGCTGTACTA-3’; Reverse: 5’-CGCTTCGGCCGGCGAAGA-3’, and *ppe11*: Forward, 5’-CGGCACCGCAAGCAACGAG-3’; Reverse: 5’-GCGGTCCCAAGTTCCCA AGT-3’. For IFN-β analysis, Forward, 5’-TCCGAGCAGAGATC TTCAGGAA-3’; Reverse: 5’-TGCAACCACCACTCATTCTGAG-3’, cDNA was synthesized using AMV Reverse Transcriptase and Oligo(dT)_20_ primer. Quantitative PCR was performed using PerfeCTa SYBR^®^ Green SuperMix (Cat. 95054, Quantabio) and specific primers on StepOnePlus Real-Time PCR System (Applied Biosystems). GAPDH: Forward, 5’-TCGTCCCGTAGACA AAATGG-3’; Reverse: 5’-TTGAGGTCAATGAAGGGGTC-3’, was used as an input control.

### Liposome RNA Treatment

EV RNA from WT *M.tb*-infected BMMs was prepared as described above using mirVana™ miRNA Isolation Kit (AM1560, Invitrogen) and purified RNA was packed using Lipofectamine 2000 (Cat.11668-027, Invitrogen) following the manufacturer’s protocol.

WT BMMs were treated with EVs (10 μg/ml) from macrophages infected with WT, Δ*secA2* or *secA2* complementary strains in the presence of liposome RNA at a final concentration of 10 pg/ml when required. xcfv

### Whole-cell Lysates and Nuclear Fraction Preparation

BMMs were treated with EVs from uninfected or *M.tb*-infected macrophages at a dose of 10 μg/ml for 4 hr at 37ºC and 5% CO_2_, and then washed with pre-cold PBS three times. Whole-cell lysates (WCL) were prepared by adding WCL lysis buffer (50 mM Tris-HCl, pH 8.0, 150 mM NaCl, 1.0% Triton X-100) containing 1x protease inhibitor cocktail (P8340, Sigma-Aldrich) and incubated on ice 30min. Nuclear Fraction was prepared as described previously (Flaherty et al., 2002). Briefly, cells were washed with pre-cold PBS three times and lysis buffer (10 mM HEPES (pH 7.8), 10 mM KCl, 2 mM MgCl2, 0.1 mM EDTA) once. Cell pellets were then resuspended in 500 μl of lysis buffer containing 1x protease inhibitor cocktail and incubated in ice for 15 min before adding 25 μl of 10% NP-40 and mixed thoroughly. Cell lysates were centrifuged at 1,200 xg for 10 minutes at 4°C. The pellets (Nuclear Fraction) were washed with lysis buffer three times and resuspended in WCL lysis buffer and incubated on ice for 30min before adding SDS-loading buffer (5x).

### Immunoblotting

WCL and nuclear fraction were denatured at 95°C for 10 min and separated by 12.0 % SDS-PAGE gel. Proteins were transferred onto nitrocellulose membranes and probed with rabbit anti-IRF3 (Cat.A303-384A, Bethyl Laboratories Inc), anti-TBK1 (Cat.3504, Cell Signaling Technology), anti-phospho-TBK1 (Ser172) (Cat.5483, Cell Signaling Technology), anti-β-actin (Cat.4970, Cell Signaling Technology), and anti-Histone H3 (Cat.9717, Cell Signaling Technology) antibodies, followed by goat anti-rabbit IgG-HRP (Cat.31460, Thermo Scientific).

### Immunofluorescent Microscopy Analysis

For EVs pretreatment assay, mouse BMMs (1 × 10^5^ cell/well) were seeded onto glass coverslips overnight, and then pretreated with 10 μg/ml EVs for 5 hr before infected by *M.tb* strains at an MOI of 5 at 37°C and 5% CO_2_ for 1 hr, followed by three washes with complete BMM medium. The infected cells were incubated at 37°C and 5% CO_2_ for another 24 hr, and fixed in 4% paraformaldehyde (PFA) at RT for 2 hr. The fixed cells were permeabilized in PBS containing 0.2% Triton x-100, and then blocked in PBS plus 2% FBS for 30 min at RT and incubated in primary rabbit anti-Lamp-1 (Cat.No. sc-5570, Santa Cruz Biotechnology, Inc.), anti-LC3 (Cat.No.12741, Cell Signaling Technology), anti-NOX2 (Cat.No. 611414, BD), anti-p47^phox^ (Cat.No. SC-17844, Santa Cruz Biotechnology) or Mono-and polyubiquitinylated conjugates monoclonal antibody (FK2) (Cat.No. BML-PW8810-0100, Enzo Life Sciences) antibody for 1hr at RT. The cells were then washed with PBS three times and incubated with Cy3-conjugated donkey anti-rabbit (Cat.No. 711-165-152, Jackson ImmunoResearch) or Texas Red-conjugated goat anti-mouse (Cat.No. T-6390, Invitrogen) IgG secondary antibody for 1 hr at RT. The coverslips were washed in PBS three times and mounted onto the glass slides. For post-exposure treatment, BMMs were infected with *M.tb* at an MOI of 5 at 37°C and 5% CO_2_ for 1 hr before being washed with complete BMM medium three times. After 24 hr, *M.tb*-infected cells were treated with EVs (5μg/ml), moxifloxacin (1.0 μg/ml) or combination for another 24 hr before immunostaining. To determine IRF3 localization in the nucleus, BMMs cells were treated with EVs at 37°C and 5% CO_2_ for 4 hr and then probed using rabbit anti-IRF3 antibody and cy3-conjugated goat anti-rabbit IgG secondary antibody as described above. The nuclei of cells were stained with DAPI. The slides were analyzed using Nikon C2^+^ Confocal laser scanning microscope at Optical Microscopy Core, University of Notre Dame. For quantitative analysis, at least 100 cells every condition were counted in three independent areas of slides.

### Combination Treatment with EVs and Moxifloxacin in *M.tb*-infected Mice

*Mavs*^−/−^ and wild-type C57BL/6 mice (8-10 weeks old, female) were infected with wild-type *M.tb* H37Rv via a low-dose aerosol infection, 100-150 CFUs in the lung, using a Glas-Col Inhalation Exposure System (Glas-Col, Terre haute, IN) as described previously (Cheng et al., 2014). *M.tb* input in the lung of mice was determined at day 1. Three weeks post-infection, infected mice were treated by oral gavage with Moxifloxacin at a dose of 50 mg/kg daily six days per week for 2 weeks, and one dose of EVs (5 μg/mouse, in 50μl PBS) was administered intratracheally at 4 week post *M.tb* infection as described previously (Cheng et al., 2017). After the treatment, mouse serum was harvested via cardiac puncture and prepared using BD Microtainer Serum Separator Tube (BD), and cytokine production was measured by ELISA as described below. In parallel, mouse lungs and spleens were harvested, homogenized, and plated onto Middlebrook 7H11 agar plates, and mycobacterial colonies were counted after 3-4 weeks of incubation at 37 °C and expressed as log10 CFUs per organ. For pathological analysis, mouse lung sections were prepared and stained with hematoxylin and eosin (H&E) at the Histology Core Facility of University of Notre Dame, and scored as described previously (Cheng et al., 2014). *M.tb* infections were carried out in the biosafety level 3 laboratory.

### ELISA

IFN-β and TNF-α level was measured in BMMs culture supernatants 24 hr after EV treatments. ELISA was performed according to the manufacturer’s instruction (ebioscience). Avidin-HRP (Cat.18-4100-94, ebioscience), and TMB (Cat.00-4201-56, ebioscience). For TNF-α, capture antibody (Cat.14-7341-85, ebioscience), detection antibody (Cat.13-7341-85, ebioscience), TNF-α Standard (Cat.39-8321-60, ebioscience). For IFN-β, capture antibody (Purified anti-mouse IFN-β Antibody, Cat. 519202, Biolegend), detection antibody (Biotin anti-mouse IFN-β Antibody, Cat. 508105, Biolegend), IFN-β standard (Cat. 581309, Biolegend). Mouse IFN-γ and IL-1β was measured using IFN gamma (Cat.88-7314-22, ebioscience) and IL-1 beta (Cat.88-7013-22, ebioscience) mouse ELISA kit, respectively.

### Statistical Analysis

Statistical analysis was performed to determine differences between groups by two-tailed Student’s t-tests using GraphPad Prism software (Version 5.04, Graphpad Software). *P*-value < 0.05 was considered significant.

## Acknowledgments

Funds for this work were provided by the grant AI052439 from the National Institute of Allergy and Infectious Diseases. We are deeply grateful to Dr. Stanley Perlman from University of Iowa for providing the MAVS^−/−^ mice. We are also very appreciative to the TARGET Program at the John Hopkins University School of Medicine for providing us the SecA2 CDC1551 mutant.

## Author Contribution

Y. Cheng was responsible for the experiments performed in this study and assisted in the writing of the manuscript. J. Schorey aided in the design of the experiments and wrote the manuscript.

